# Comparing two automated methods of DNA extraction from degraded skeletal remains

**DOI:** 10.1101/2020.09.22.308858

**Authors:** Linda Rubinstein

**Affiliations:** NASA

## Abstract

DNA extraction from degraded skeletal samples is often particularly challenging. The difficulty derives from the fact that variable environment has significant effect on DNA preservation. During the years 2002-2015 unidentified degraded skeletal remains were accumulated at our institute, National Institute of Forensic Medicine (NIFM), most of them with none or partial DNA profile.

As new methods rapidly emerge, we revisited the samples with partial DNA profiles. We have chosen to use these samples to compare two automated methods: Prepfiler Express BTA (Applied Biosystems) and QIAcube (Quiagen), in hope of acquiring a more complete DNA profile and eventually even make new identifications. In both methods a preparation step is required, after which the samples undergo automatic DNA extraction.

The two protocols are based on different extraction methods. Fresh or non-problematic bone samples as the positive control gave the same results in both methods. In the degraded skeletal samples, the results were significantly better using the QIAcube method.

## Introduction

Extraction of DNA from degraded skeletal remains continues to pose a great challenge for forensic scientists worldwide. The need for this kind of extraction may occur in cases of mass disasters, forensic criminal casework, missing persons cases or historical war victims [1]. The human tissue is rapidly degraded and in many identification cases the only possibility of obtaining DNA is from bone or teeth. There are many factors affecting the levels of sample preservation: surface exposure to UV, heat, aquatic environment, microbial activity and more [2, 3].

As degradation proceeds, DNA becomes more fragmented and successful typing of STR profiles is decreased. Determining of Mitochondrial DNA (mtDNA) is a quite reliable mean of identification but the statistical power generated by this method is relatively low [4]. In addition to the DNA being degraded the samples may also be structurally damaged including DNA-DNA cross linkage or contamination. The DNA extraction methods are improving lately and even being automated, but still there are many challenges left.

In this study we aimed to compare two semi-automated extraction methods - AutoMate Express (Applied Biosystems) using PrepFiler™ Forensic DNA Extraction Kit and the QIAcube (Quiagen) using QIAquick^®^ PCR Purification Kit, in highly degraded skeletal samples.

PrepFiler™ BTA Forensic DNA Extraction Kit according to the manufacturer, is specifically designed to improve the yield and purity of DNA prepared from forensic samples. The method involves binding DNA to coated magnetic particles in the presence of chaotropic salts, washing of the particles to remove undesirable compounds, and elution of DNA from the particles in a low-salt solution (1).
QIA Cube^®^ according to the manufacturer, is an extraction protocol that combines the “full demineralization” process of the bone according to Loreille *et al*, [4] with the silica based cleanup of Yang *et al* [7], it is automated using the Qiagen QIAcube automated sample preparation instrument as described in Amory *et al* [5].

## Materials and Methods

### Sample Preparation

Twenty-six samples taken from the workflow of our lab were chosen from degraded skeletal remains - most of them from sculls.

The preparation and cleaning of the remains was the same in both methods: The bone/scull was cleaned with DDW, dried and the surface removed using a sanding machine (Horico), to eliminate potential contamination.

Following this cleaning procedure, the samples were ground in the presence of liquid nitrogen. From each sample 0.5mg of bone powder was submitted for extraction.

### DNA extraction methods

#### AutoMate Express (Applied Biosystems)

The bone powder was incubated overnight in 0.5M EDTA (Amnion) at 37°C on a rocking platform. The next day the samples were centrifuged and transferred into 2ml tubes. The pellet was rinsed twice in DDW and then lysis buffer from the Prepfiler BTA kit (PrepFiler^®^ Express BTA Forensic Extraction Kit (Applied Biosystems) was added together with DTT (Sigma) and PK (Roche) as recommended. The samples were than incubated overnight in thermos-shaker on a rocking platform at 950 rpm at 56°C. The next day the samples were submitted to the automatic robot extraction as previously described [6].

#### QIAcube (Quiagen)

The bone powder was incubated overnight in 0.5M EDTA and 500μl of proteinase k 20mg/ml (Roche), vortexed and incubated ON in a thermo shaker at 56°C. The following day the samples were centrifuged for 5 minutes at 1800Xg. The supernatant was transferred to an Amicon Ultra^®^ 15 ml – 100kDA. The Amicon column was centrifuged at 1800Xg till the samples were concentrated to a 300 μl volume. The recovered solution was transferred from the Amicon filter to a 2 ml tube and the samples 1500 μl of Buffer PBI (QIAquick^®^ PCR Purification Kit) was added and the samples were purified as instructed by the QI manual were submitted to the automatic robot extraction as previously described [8].

#### DNA amplification, Fragment separation and data analysis

All DNA extracts were amplified using the Powerplex ESI/ESX 16 System (PP16, Promega).

Fragment separation was performed using AB Hitachi 3500XL genetic analyzer using POP-4 polymer and the version 1.2 GeneMapper^®^ ID-X Software. (Applied Biosystems). All profiles were compared to the laboratory staff database to ensure lack of contamination.

### Results and Discussion

We aimed to extract DNA from degraded skeletal remains of unknown persons found in Israel during the years 2002-2015 and compare two distinct DNA extraction methods. We didn’t chose the samples specifically we have just used samples which in the past didn’t give sufficient results hoping that the new methods would improve the allele detection and by this enabling better chances of the sample being found in the missing persons database. Many of the samples found in those years were from skulls probably because they are preserved better in terrain than other bones. We have compared two semi-automated extraction methods QIA Cube and AutoMate Express. In both methods most of the profiles obtained were still partial since most of the samples were degraded. The level of detection was set above 100 RFU. Reliable result based of at least allele duplication. Table1 summarizes all the samples in our study, including the environment and the year in which they were found.

The QIAcube method showed statistically better results in the degraded samples (Figure 1). We did not observe any correlation between the environments the samples were found in and the differences between the two methods. It would be interesting to compare these methods with more samples from different terrains (water, soil, dessert, caves) since this could have additional effect on the DNA determination.

**Figure 1:**
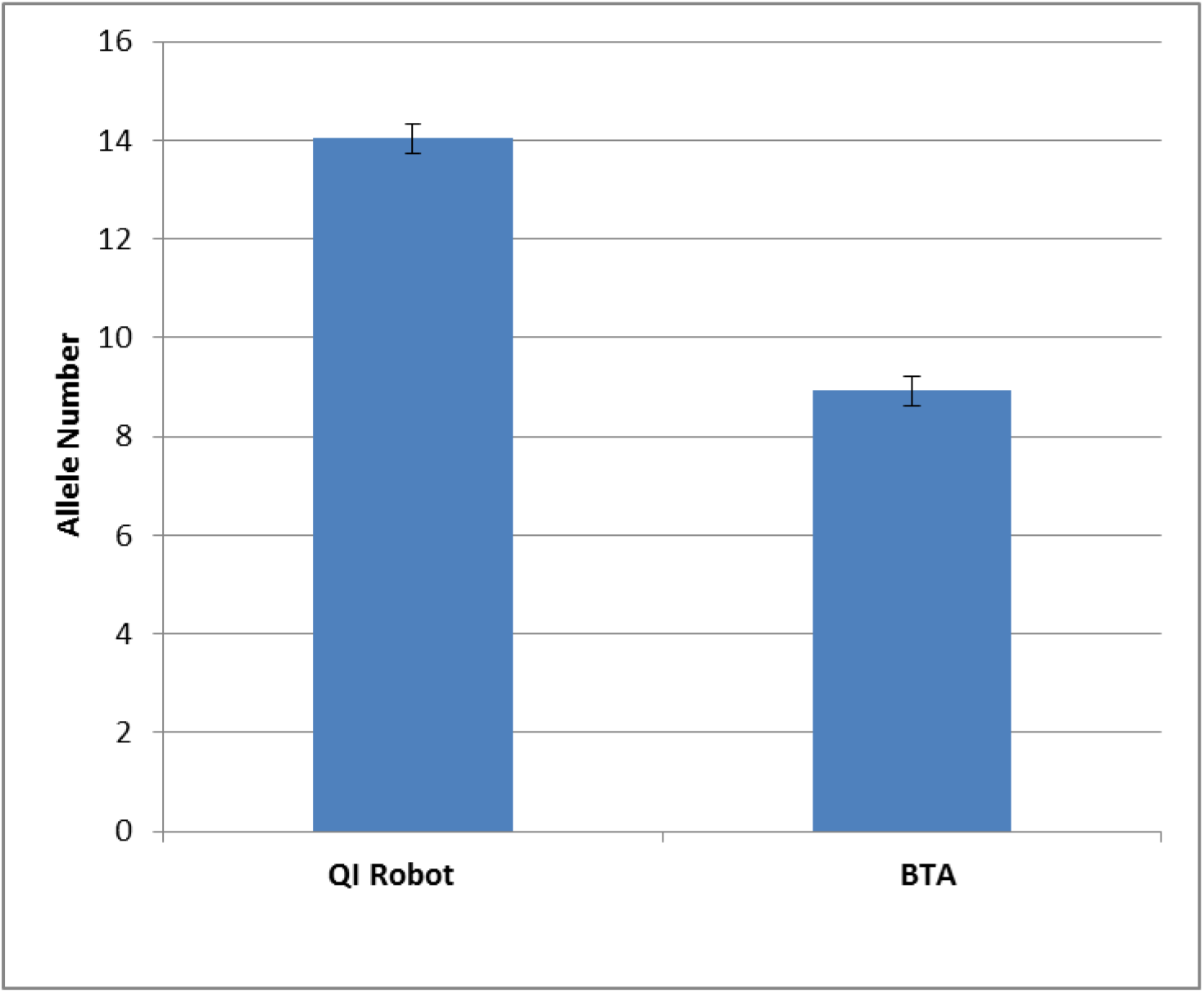
Comparing the number of alleles obtained from two automated DNA extraction methods; P<0.02.

We obtained full profiles, by the QIAcube method, even in cases in which DNA detection has failed before, however this method was not better in **all** the samples. Both methods use chaotropic salts in the environment of the DNA extraction. The main difference between the two methods is that QIAcube uses silica columns while the AutoMate Express uses magnetic beads for the extraction. The capacity of the silica columns is 10 μg and the capacity of the magnetic beads is 2μg.

We hypothesize that in some degraded samples the more limited capacity of binding DNA to the beads may affect the final yield. In other samples affected by the microorganisms and environment the purity of the final product submitted to PCR will be more important. Practical considerations using these two methods include: time, amount of samples and possibility of contamination. More samples can be extracted simultaneously in the Prepfiler than the QIAcube but since it is not recommended to extract more than one bone simultaneously due to possible cross contamination this is not a problem. The time of extraction is similar in both methods. The additional time-consuming step in the QIAcube method is the Amicon filter centrifugation, in which some difficult samples may take up to one hour, but overall, the automation process takes about the same time. The automation process itself is has an advantage in avoiding contamination as fewer manual steps and handling are required.

From this work it seems that the QIAqube method is more suitable for degraded samples (Figure 1) but since in some samples we received better results in the PrepFiler method and since degraded samples are highly variable we would recommend to use both methods in case of difficult samples and by this combination receive better and more reliable profiles.

In future studies it would be also interesting to compare these two methods in remains from various anatomical parts of the skeleton. For example results in DNA extraction from the petrous part of the temporal bone had shown good results [7].

Automation of the forensic casework and identification cases is important to achieve more reliable, rapid and high-throughput sample processing. We hope that the results of this study will contribute to forensic cases dealing with skeletal remains.

**Table1:** The number of alleles obtained by the QIA Cube and the AutoMate Express methods, the sample specifications and year of finding.

| **Sample**<br>**Number** | **QIA**<br>**Cube**<br>**Number**<br>**of alleles**<br>**above**<br>**100 RFU** | **AutoMate**<br>**Express**<br>**Number**<br>**of alleles**<br>**above 100**<br>**RFU** | **Details and Year in which the remains were found** |
| --- | --- | --- | --- |
| 1 | 32 | 26 | Human skull found at seashore (2013) |
| 2 | 10 | 18 | Mandible bone found at seashore (2015) |
| 3 | 8 | 8 | Femur buried in the ground (2013) |
| 4 | 5 | 10 | Teeth (2015) |
| 5 | 24 | 30 | Human skull found in a cave (2015) |
| 6 | 2 | 2 | Human skull found in a cave (2011) |
| 7 | 4 | 2 | Human skull (2008) |
| 8 | 28 | 13 | Human skull buried in the ground (2004) |
| 9 | 24 | 14 | Part of human bone found in open area (2012) |
| 10 | 21 | 21 | Mandible bone found in a cave (2009) |
| 11 | 31 | 1 | Human skull found at seashore (2013) |
| 12 | 10 | 0 | Part of human bone found in open desert area (2014) |
| 13 | 8 | 0 | Human skull found in the seawater (2013) |
| 14 | 4 | 6 | Human skull found inside a building (2015) |
| 15 | 7 | 1 | Human skull (2011) |
| 16 | 2 | 0 | Human skull found in fresh water (2009) |
| 17 | 1 | 0 | Human skull found in the seawater (2014) |
| 18 | 5 | 0 | Parts of human skeletal remains burned (2010) |
| 19 | 17 | 24 | Human skull found in open area (2006) |
| 20 | 25 | 16 | Human skull found in desert area (2005) |
| 21 | 22 | 25 | Human skull found in open area (2002) |
| 22 | 14 | 0 | Parts of human skeletal remains found in open area (2007) |
| 23 | 24 | 3 | Part of human skull found in forest area (2008) |
| 24 | 10 | 0 | Human skull (2009) |
| 25 | 13 | 3 | Human skull (2008) |
| Positive Control | 30 | 30 |  |

